# Heterogeneous T cell responses across Hepatitis B virus clinical phases revealed by rapid whole-blood HBV T cell analysis

**DOI:** 10.1101/2025.02.11.637593

**Authors:** Nina Le Bert, Apostolos Koffas, Rajneesh Kumar, Floriana Facchetti, Anthony T Tan, Upkar S Gill, Lung-Yi Mak, Sophie W Stretch, Tizong Miao, Smrithi Hariharaputran, Shou Kit Hang, Yifei Guo, Qi Chen, Elisabetta Degasperi, Pietro Lampertico, Wan Cheng Chow, Antonio Bertoletti, Patrick TF Kennedy

**Affiliations:** Emerging Infectious Diseases, Duke-NUS Medical School, Singapore; Barts Liver Center, Barts and The London School of Medicine & Dentistry, Queen Mary University of London, London, UK; Department of Gastroenterology and Hepatology, Singapore General Hospital, Singapore; Gastroenterology and Hepatology Division, Foundation IRCCS Ca’ Granda Ospedale Maggiore Policlinico, Italy; T Cell Diagnostics Ptd. Ltd., Singapore; CRC “A. M. and A. Migliavacca” Center for Liver Disease, Department of Pathophysiology and Transplantation, University of Milan, Milan, Italy

**Author notes:** **Authorship Contributions**: Conception/design: NLB, AB, PTFK; Patients selection: AK, RJ, ED, PL, WCC, PTFK; Data collection and/or processing: FF, USG, LYM, SWS, TM, SH, SKH, YG, QC; Analysis and Interpretation: NLB, ATT, PL, AB, PTFK; Visualization: NLB, ATT; Manuscript draft: NLB, AB, PTFK; Manuscript editing: ATT, ED, PL, WCC; Manuscript review: all authors.

**Keywords:** Hepatitis B virus, CHB, T cells, cytokines, liver disease, biomarker

## Abstract

**Background & Aims:** There is an unmet need for immunological biomarkers in chronic hepatitis B (CHB), where patient management relies on virological and biochemical markers despite the crucial role of virus-specific T cells in controlling viral replication and disease progression. To address this, we developed the HBV-cytokine release assay (HBV-CRA), a rapid, point-of-care test that measures multiple cytokines in whole blood after HBV-peptide stimulation.

**Methods:** We first assessed the assay’s sensitivity by spiking whole blood with engineered HBV-specific T cells. Next, we compared sensitivity and consistency of HBV-CRA to ex vivo IFN-γ ELISpot assays. We then applied the assay in a cross-sectional study of 235 CHB patients and longitudinally in acute HBV patients during HBsAg sero-clearance.

**Results:** The HBV-CRA detected T cell function in 80% of CHB cases and showed that elevated IL-2 and IFN-γ levels after Core peptide stimulation were associated with HBsAg clearance. Importantly, unsupervised clustering identified distinct immune response patterns independent of established clinical and virological classifications. The assay also demonstrated the functional impact of NUC treatment on HBV-specific T cell responses.

**Conclusions:** The HBV-CRA is a rapid and easy-to-use assay that identifies immune profiles associated with HBsAg clearance and differentiates CHB patients based on antiviral T cell function. Its application in a large CHB cohort revealed that traditional disease phase classifications, based on viral and clinical parameters, cannot predict HBV-specific T cell profiles. The HBV-CRA has the potential to guide patient stratification for immunotherapeutic interventions.

## Introduction

Functional and quantitative defects of HBV-specific T cells are likely the primary reason chronic HBV infection (CHB) patients cannot achieve HBsAg seroconversion(1). Research showed that a high frequency of HBV-specific CD8+ T cells in the bloodstream, observed only in acute but not chronic HBV infection, leads to HBsAg loss and a 90% reduction in HBV-DNA levels(2). Furthermore, while acute HBV patients possess functional HBV-specific Th1-like CD4 T cells, CHB patients exhibit dysfunctional HBV-specific CD4 T cells(3,4), which likely contribute to the failure of HBs-specific B cells to fully mature and produce anti-HBs antibodies(5,6).

While acute HBV patients show a consistent profile of robust antiviral immunity, the adaptive immune defects in CHB patients are highly variable. HBV-specific T cells in CHB are present at low frequencies (<0.1% of T cells)(7–10) and show heterogeneity in exhaustion marker expression, including PD-1, TIM-3, Lag-3, and CD160(10–14). Functionally, these T cells exhibit impaired proliferation and cytokine production(4,8,12,15). Their defects are heterogeneous(16), not exclusively associated with the expression level of inhibitory receptors(14) and not solely related to classical T cell exhaustion mechanisms due to persistent antigenic stimulation(17). These defects vary with HBV antigen specificity(10,11,16,18), interaction with liver endothelial cells(19) and their “priming history”, as T cells primed by hepatocytes or dendritic cells possess distinct functional profiles(20). Although better HBV-specific T cell function can sometimes be observed in CHB patients with low viral load(11,14,18) or younger age(21), we cannot interpret the intricate spectrum of HBV-specific T cells through the standard virological and clinical assessments like HBsAg, HBeAg status or ALT levels. Only advanced immunological methods can define the complex nature of the HBV-specific T cells in CHB patients(22). Still, these methods are limited to highly specialized laboratories and not easily translated into clinical practice for large clinical trials.

Thus, we developed a simple, point-of-care immunological test that can define the functional profile of HBV-specific T cells of acute HBV patients during the dynamic phase of HBV control and the complex heterogeneity of HBV-specific T cells in CHB patients.

We modified a method that we developed to rapidly measure the frequency and function of SARS-CoV-2-specific T cells(23,24). We designed HBV peptide pools covering multiple genotypes and used them directly in the blood of acute HBV and CHB patients to stimulate the release of cytokines from HBV-specific T cells. After overnight incubation, IFN-γ, IL-2, IL-10, granzyme B (GrzB), TNF-α, and IL-5 were measured. We demonstrated that this simple HBV-cytokine release assay (HBV-CRA) enables the quantitative and functional profile of HBV-specific T cells using minimal blood volume. By applying the HBV-CRA in a large cohort of 235 CHB patients, we identified distinct HBV-specific T cell functional profiles, or secretomes, even among patients classified within the same clinical and virological categories, allowing their immune-based stratification.

## Material and Methods

### Study approval

Clinical assessment and blood sampling were performed at The Royal London Hospital, Barts Health NHS Trust, Singapore General Hospital and Policlinico of Milan in accordance with the Declaration of Helsinki. Written informed consent was obtained, and the Institutional Review Boards approved the study (refs: 17/LO/0266, 2022/2732, ID-4670).

### Blood samples

260 individuals were included in this study: HBV-vaccinated subjects (n=16), treatment-naïve CHB patients (n=100), NUC-suppressed CHB patients (n=74), HBsAg-negative CHB patients (n=61), and patients with acute resolving HBV infection (n=9). A subset of NUC-suppressed patients (n=10) was sampled monthly over three time points. Acute HBV patients were tested 1-4 times during viral clearance. Tables S1, S2 and S3 summarized the clinical and virological parameters.

### Peptide pools

We used a library of 838 overlapping 15-mer peptides (10-aa overlap). These peptides were combined into six pan-genotype pools covering all HBV proteins from genotypes A, B, C, and D and sourced from T Cell Diagnostics. SARS-CoV-2-Spike peptide pool was generated as described before(23).

### Ex-vivo ELISpot assays

PBMCs were seeded at 400,000 cells per well onto anti-IFN-γ (1-D1K, Mabtech; 5 μg/mL) pre-coated 96-well plates. Cells were stimulated for 18h with peptide pools (2 μg/mL), DMSO, or PHA controls. Plates were developed using biotinylated IFN-γ detection antibody (7-B6-1, Mabtech; 1:2000), Streptavidin-AP (Mabtech), and KPL BCIP/NBT substrate (SeraCare). We quantified Spot-forming cells (SFC) with ImmunoSpot, and positive responses were calculated by subtracting 2x SFC of DMSO controls from peptide-stimulated wells. All controls met quality criteria, with <5 SFC in negative controls and positive PHA controls.

### HBV-Env_183-91_ TCR T cells

HLA-A2-restricted HBV-Env_183-91_-specific T cells were generated by electroporating mRNA encoding the TCR Vα and Vβ chains into 10-day expanded T cells from an HLA-A2+ subject(25). HBV-Env_183-91_-TCR expressing T cells were quantified via dextramer staining.

### Whole blood HBV-CRA

On the same day of collection, whole blood drawn into heparinized tubes (320 μL blood per peptide pool) was mixed with 80 μL RPMI and stimulated with HBV peptide pools (2 μg/mL) or DMSO control. After 16 hours of culture at 37°C, the culture supernatant (plasma) was collected and stored at -80°C for cytokine quantification.

### Cytokine quantification and analysis

We measured IFN-γ, IL-2, TNF-α, IL-4, IL-5, IL-10, and GrzB in the plasma using an Ella machine (ProteinSimple). Cytokine levels in DMSO controls were 2x subtracted from peptide-stimulated samples. Dimensionality reduction was performed using UMAP(26) and Phenograph(27) with nearest neighbors = 200, min_dist = 0.99, k = 30. UMAP results were converted to .fcs files and analyzed in FlowJo to generate cytokine secretion heatmaps.

### Statistical analysis

All tests are stated in the figure legends. P-values (all two-tailed): <0.05 = *; <0.01 = **; <0.001 = ***; <0.0001 = ****.

## Results

### Sensitivity of the HBV-cytokine release assay (HBV-CRA)

During the COVID-19 pandemic, we demonstrated that whole blood stimulation with peptide pools covering SARS-CoV-2 proteins can detect SARS-CoV-2-specific T cells more accurately than other assays (ELISpot, AIM assay)(23). However, while vaccine- or infection-induced SARS-CoV-2-specific T cells often comprise around 0.1% of total T cells, circulating HBV-specific T cells in CHB patients rarely reach this level. This different magnitude is depicted in Figure 1A, where we tested T cells of COVID-19-vaccinated CHB patients against SARS-CoV-2 Spike and HBV-proteins. While the ex vivo IFN-γ ELISpot assay detected SARS-CoV-2-specific T cells, HBV-specific T cells were rarely found. To assess whether the whole blood HBV-CRA can achieve a sensitivity threshold suitable for detecting the low frequency of HBV-specific T cells, we diluted graded numbers of T cells with known specificity into whole blood. We engineered a T cell receptor (TCR) specific for the HLA-A2-restricted HBV-Env_183-91_ epitope into T cells of an HLA-A2+ subject and we diluted decreasing numbers of HBV-Env_183-91_-TCR expressing T cells into the whole blood from the same subject (Figure 1B). After overnight culture with the corresponding HBV-Env-_183-91_ peptide, plasma cytokines were measured. Figure 1C shows that IFN-γ and GrzB levels >10 pg/mL were detected when 50-100 Env_183-91_-TCR+ T cells (∼0.01-0.02% of total T cells) were diluted in blood, matching the frequency of circulating HBV-specific T cells in most CHB patients. IL-2 was first detected at ∼100 cells/320 μL of blood, but levels >10 pg/mL were observed only at higher T cell numbers (∼1000), in line with the effector profile of these engineered TCR-T cells.

**Fig. 1.**
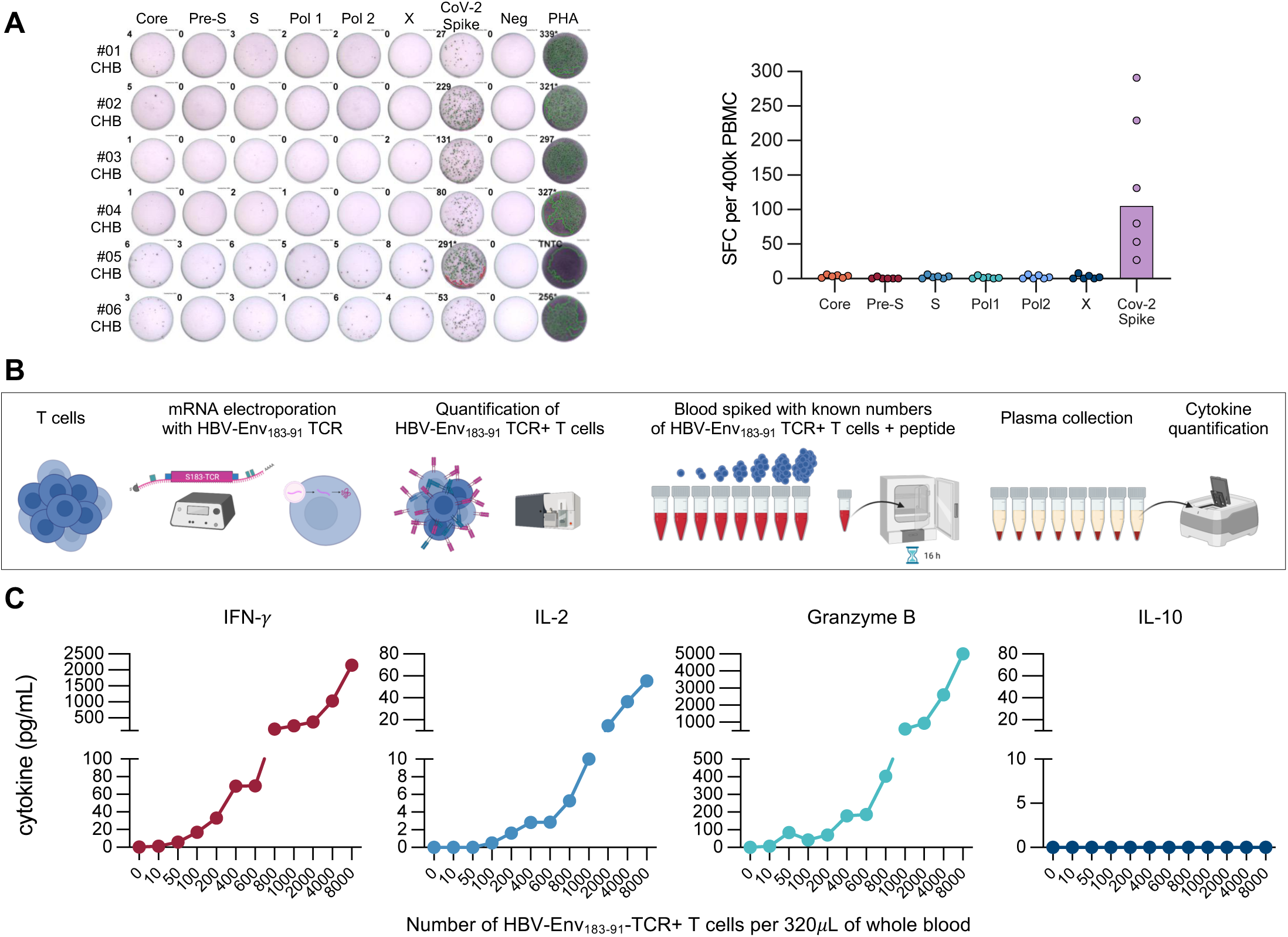
Sensitivity of the whole blood HBV-cytokine release assay. (**A**) PBMCs were stimulated with peptide pools covering the indicated HBV proteins and SARS-CoV-2 Spike. IFN-γ-spot forming cells (SFC) were quantified by ELISpot assay. (**B**) Graded numbers of HBV-Env_183-91_-TCR+ T cells were spiked into 320 μL of fresh blood of the same subject and stimulated overnight with the Env_183-91_ peptide or corresponding DMSO controls. Subsequently, cytokines were quantified in supernatants. (**C**) IFN-γ, IL-2, GrzB, and IL-10 levels induced by peptide stimulation in blood samples containing the specified numbers of HBV-Env_183-91_-TCR+ T cells.

### HBV-CRA in CHB patients and healthy controls: cytokine selection

Since the sensitivity of the HBV-CRA aligns with the known frequency of HBV-specific T cells in CHB patients, we designed six peptide pools of overlapping 15-mer peptides (10-aa overlap) covering the Core (73 peptides), S (188 peptides), PreS (35 peptides), Polymerase (split into Pol-1 with 230 peptides and Pol-2 with 231 peptides) and X (83 peptides) proteins of HBV genotypes A, B, C, and D. These were organized into individual pools (Figure 2A). Each peptide pool was added to fresh whole blood and incubated overnight to assess HBV-peptide-induced cytokine secretion. Supernatants were then analyzed for Th1 cytokines (IFN-γ, IL-2, TNF-α), Th2 cytokines (IL-5, IL-4), GrzB, and IL-10 (Figure 2B).

**Fig. 2.**
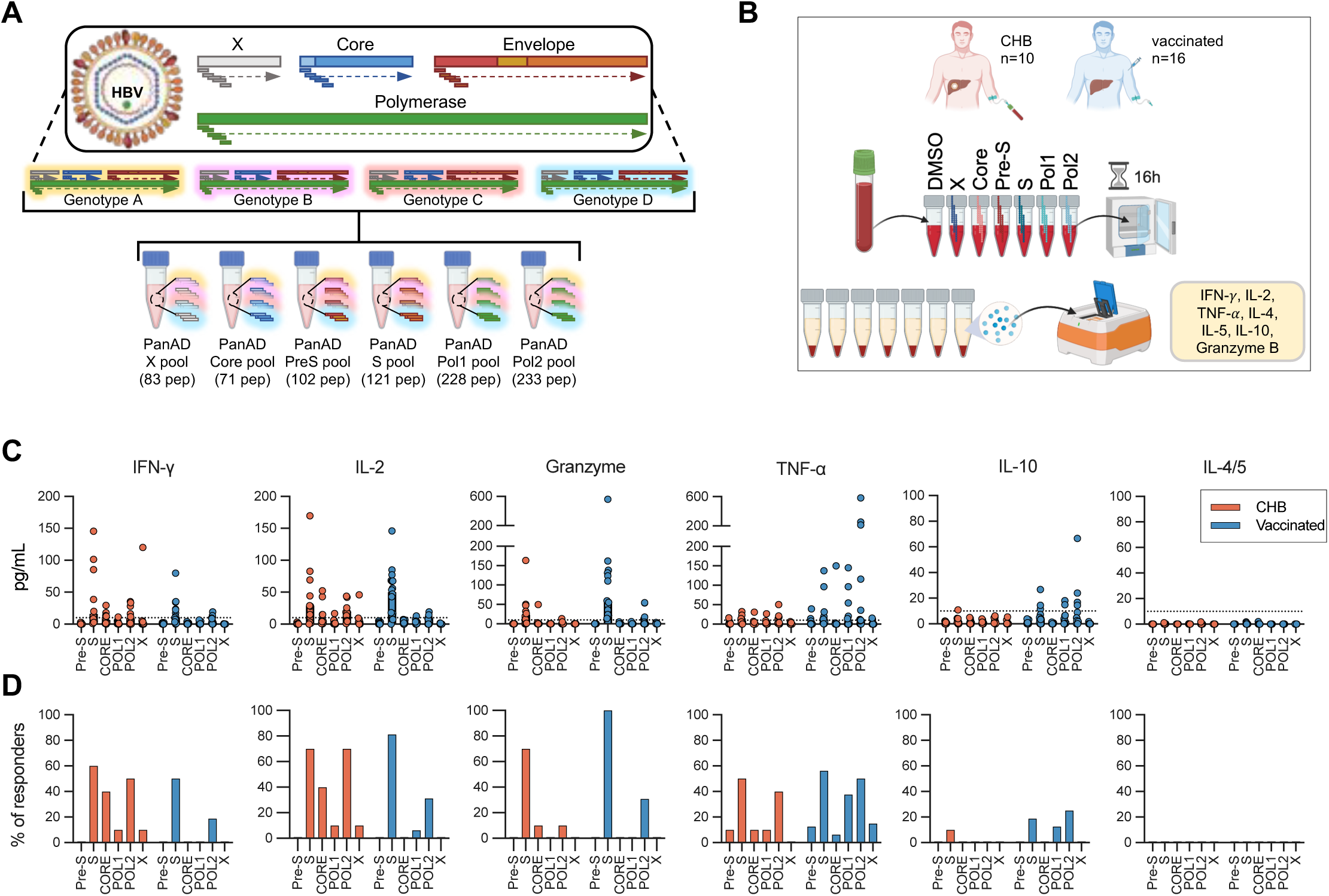
HBV-peptide induced cytokine release assay: CHB patients versus HBV vaccinated subjects. (**A**) Schematic of HBV pan-genotype (A-D) peptide pool design. (**B**) Schematic of the whole-blood HBV-CRA. (**C**) Fresh blood from NUC-suppressed CHB patients (n=10) and healthy-vaccinated subjects (n=16) was stimulated with the different HBV-peptide pools. Secreted cytokine concentrations are shown (red=CHB; blue=healthy-vaccinated). (**D**) Percentage of individuals with a peptide-induced cytokine secretion >10 pg/mL is shown (red=CHB; blue=healthy-vaccinated).

We initially tested samples from 10 virally suppressed CHB patients on NUC treatment and 16 healthy HBV-vaccinated individuals (Table S1). Figure 2C shows the cytokine concentrations in CHB patients’ and vaccinated individuals’ blood stimulated with each peptide pool. We defined positive cytokine levels exceeding 10 pg/mL after subtracting twice the background levels and the frequencies of positive responses are summarized in Figure 2D. Most CHB patients showed IFN-γ and IL-2 secretion in response to multiple peptide pools, though cytokine concentrations varied widely. Surprisingly, the S peptide pool elicited stronger IFN-γ (6/10) and IL-2 (7/10) responses in CHB patients than others. The Core pool, often considered the most T cell-immunogenic HBV protein, induced IFN-γ/IL-2 secretion in fewer patients (4/10). The Pol-2 pool, covering the C-terminal polymerase region, stimulated IFN-γ/IL-2 secretion in more than half of the patients, while Pol-1 (N-terminal polymerase) and X pools triggered responses in just one patient each. The PreS pool elicited no responses.

The cytokine response profile in vaccinated individuals differed markedly. Only the S peptide pool consistently induced IFN-γ/IL-2 secretion in most individuals, while PreS, Core, X, and Pol-1 pools did not. The Pol-2 pool induced responses in 3 of 16 individuals, confirming previous reports of polymerase-specific T cells detected via MHC pentamers in some vaccinated individuals(14,28), likely due to high HBV circulation in Asia.

GrzB secretion was predominantly detected with the S peptide pool in CHB patients and vaccinated individuals. However, while GrzB secretion always co-occurred with IFN-γ/IL-2 in vaccinated individuals, it was not always the case in CHB patients (e.g., C and Pol-2 pools).

Th2 cytokines (IL-4, IL-5) were undetectable in CHB patients and vaccinated individuals. TNF-α and IL-10 secretion were frequently observed, particularly in vaccinated individuals, often independently of IFN-γ/IL-2. This suggests they may be driven by monocyte activation rather than T cell activity, and thus, may not provide reliable information about HBV-specific T cell function.

### Enhanced sensitivity and reproducibility of HBV-CRA in NUC-suppressed CHB patients

To evaluate whether the HBV-CRA detects reproducible patterns of HBV-peptide pool-induced cytokine secretion over time, we collected blood samples from 10 CHB patients on NUCs three times at four-week intervals. Concurrently, we isolated PBMCs to conduct ex vivo ELISpot assays (Figure 3A).

**Fig. 3.**
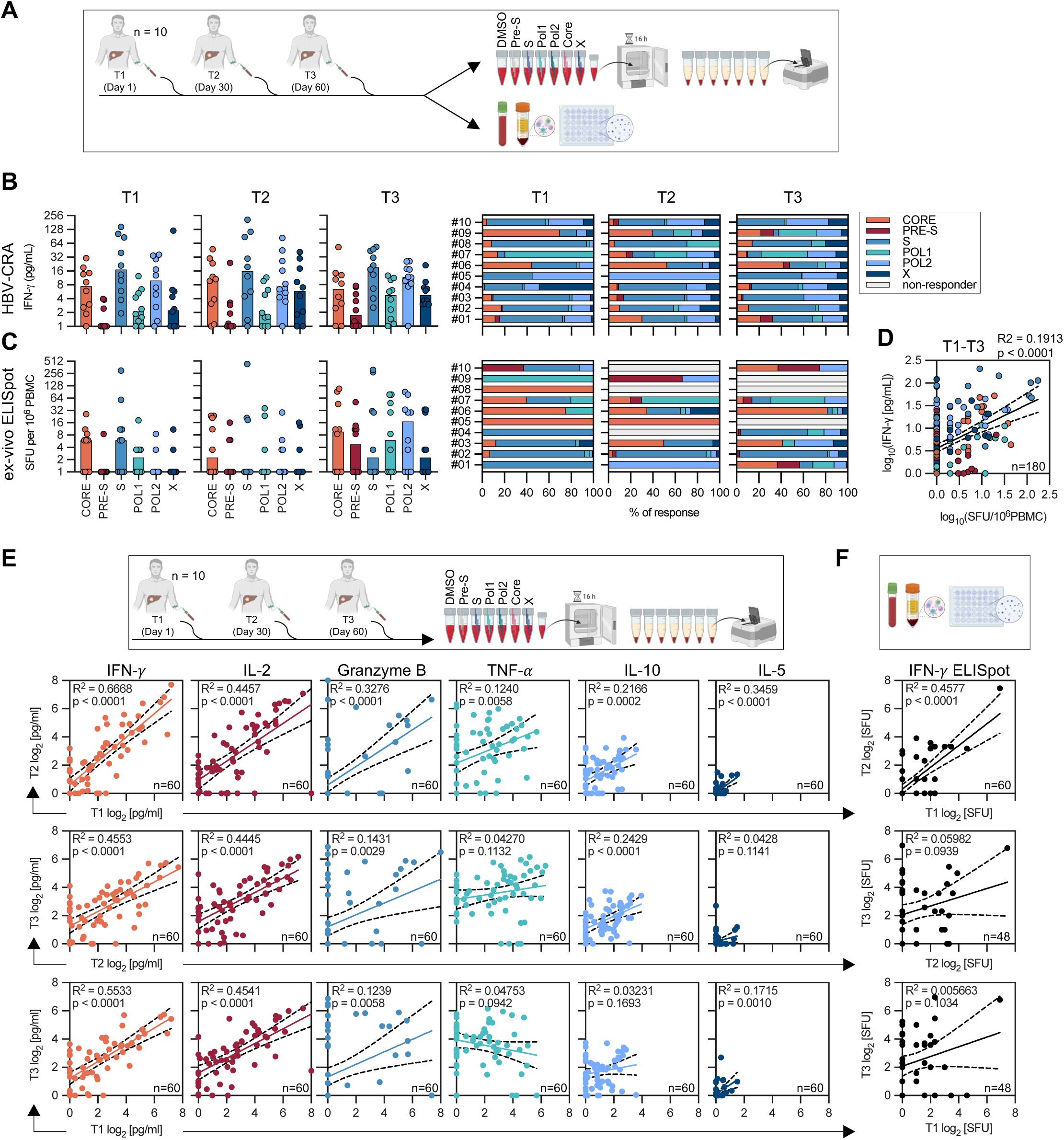
Consistency of the HBV-CRA and comparison with ELISpot assays. (**A**) NUC-treated CHB patients (n=10) were sampled monthly, and their HBV-specific T-cell profiles were analyzed with HBV-CRA and ex-vivo ELISpot assay. HBV-CRA IFN-γ-secretion levels (**B**) and ex-vivo ELISpot IFN-γ-SFUs (**C**) in response to the indicated peptide pools at T1 (Day 1), T2 (Day 30), and T3 (Day 60) (left). The percentage contribution of the single peptide pools to the total HBV-specific IFN-γ/ELISpot response is shown for individual patients across time points (right). (**D**) Correlation analysis between IFN-γ-levels from the HBV-CRA and the corresponding IFN-γ-SFUs from the ex vivo ELISpot assays in response to individual peptide pools at all time points. Correlation between (**E**) secreted cytokine levels (IFN-γ, IL-2, GrzB, TNF-α, IL-10, IL-5) and (**F**) IFN-γ-SFUs measured at different time points.

HBV-peptide-induced IFN-γ secretion showed a consistent response pattern across time points, with minor quantitative variations but stable T cell dominance in individual patients (Figure 3B). In contrast, the IFN-γ ELISpot assay exhibited greater variability, with low responses at one time point often not replicating in subsequent tests, leading to inconsistent immunodominance patterns (Figure 3C).

Correlation analysis between HBV-CRA and ex vivo ELISpot revealed that HBV-CRA often detected IFN-γ levels above 5 pg/mL without corresponding spots in ELISpot assays (Figure 3D). However, a significant correlation was found when ELISpot results exceeded 10 SFC per million PBMCs, indicating higher sensitivity of the HBV-CRA. To further evaluate the consistency of the HBV-CRA, we correlated cytokine levels at time points T1, T2 and T3 (Figure 3E). IFN-γ and IL-2 levels were highly consistent across all time points. GrzB levels also showed strong consistency between T1 and T2, though distinct responses emerged at T3, yet maintaining a significant correlation overall. TNF-α secretion was frequently detected but showed substantial variability over time, while IL-10 remained low but consistent across adjacent time points (T1 vs. T2 and T2 vs. T3). IL-5 remained consistently low or undetectable. In contrast, the correlation analysis of IFN-γ ELISpot results over time confirmed its variability, with a strong correlation only between T1 and T2.

### HBV-CRA captures dynamic kinetics of HBV-specific T cells in acute HBV infection

Building on the reproducibility of the HBV-CRA in NUC-treated CHB patients, we evaluated its ability to capture dynamic changes in HBV-specific T cell responses during acute resolving HBV infection. Unlike NUC-suppressed CHB patients, acute HBV patients are characterized by a well-documented expansion and contraction of HBV-specific CD8+ and CD4+ T cells with a Th1 cytokine profile(29–31).

We longitudinally followed nine patients with acute HBV, performing the HBV-CRA at 1-4 different time points (Table S2). Among these, three patients were sampled from the “early” acute phase (HBsAg+, HBV-DNA+, elevated ALT) to HBsAg seroconversion and ALT normalization (Figure 4, upper graphs). Cytokine secretion in response to HBV peptide pools was measured at each time point, with Core, S and Pol-2 pools stimulating responses in patient #6 and Core, S and X in patients #4 and #8.

**Fig. 4.**
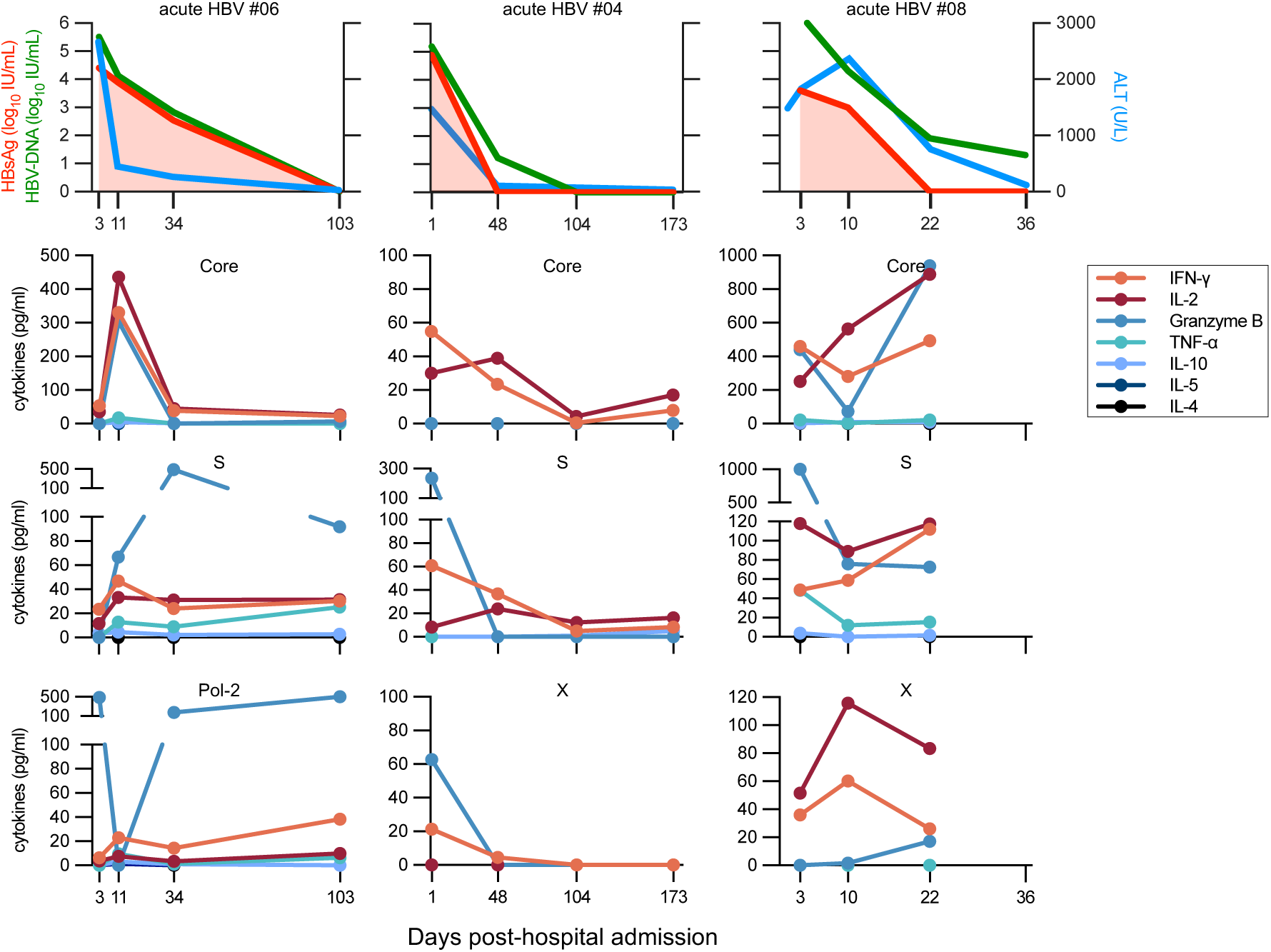
Fluctuation of the HBV-CRA in patients with acute HBV infection. Three acute HBV patients were followed longitudinally until HBsAg and HBV-DNA sero-clearance. HBV-DNA, HBsAg, and ALT levels are shown (upper panels). HBV-CRA was performed at the indicated days post-hospital admission. Cytokine secretion levels are shown in response to the indicated HBV-peptide pools.

The HBV-CRA effectively tracked dynamic HBV-specific T cell fluctuations associated with viral and HBsAg sero-clearance. Th1 cytokines (IL-2 and IFN-γ) and GrzB were consistently detected, while Th2 cytokines, TNF-α and IL-10 were minimal. IL-2 and IFN-γ showed coordinated kinetics, while GrzB followed a distinct trajectory, suggesting differential regulation.

Thus, the HBV-CRA can quantify and characterize HBV-specific T cell responses during acute infection.

### Marked heterogeneity in the HBV-specific T cell secretome among CHB patients

We then applied the HBV-CRA to a large cohort of CHB patients spanning all disease phases (Figure 5A, Table S3). This cohort included treatment-naïve (n=100) and NUC virally suppressed CHB patients (n=74), as well as 61 HBsAg-negative patients, comprising those who had either spontaneously resolved CHB infection (n=24) or achieved HBsAg sero-clearance following NUC therapy (functional cure; n=37). Additionally, we included longitudinal samples (1-3 time points, all HBV-DNA+, n=20) from 9 patients with acute HBV in the analysis.

**Fig. 5.**
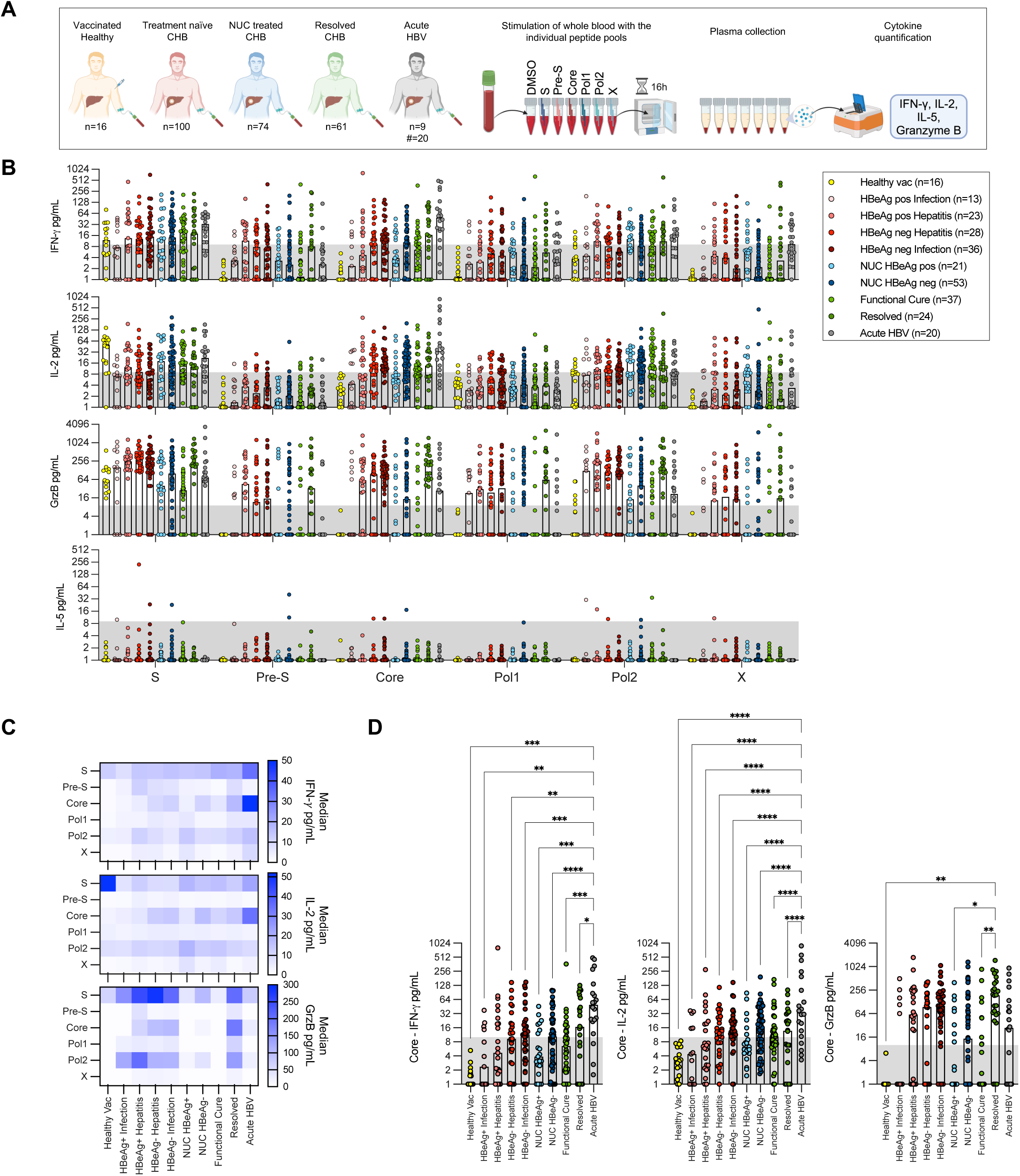
Marked heterogeneity in the HBV-specific T cell secretome among CHB patients. (**A**) Schematic of patients and HBV-CRA (n=number of subjects; #=number of samples). (**B**) Secretion levels of IFN-γ, IL-2, GrzB, and IL-5 in response to the indicated HBV-peptide pools. Each dot represents one patient sample; bar represents median. (**C**) Heatmap showing median secretion levels of IFN-γ, IL-2, GrzB in response to the six HBV-peptide pools for each patient group. (**D**) IFN-γ, IL-2, GrzB secretion levels in response to the HBV-Core peptide pool for the various patient groups. Each dot represents one patient sample; bar represents median. Ordinary one-way ANOVA followed by Tukey’s multiple comparisons test.

First, cytokine secretion levels (IFN-γ, IL-2, GrzB, IL-5) in response to each HBV peptide pool were analyzed across all samples, excluding TNF-α and IL-10 due to their likely monocyte-driven production. We observed considerable heterogeneity in cytokine secretion levels within all patient groups (Figure 5B), with the S, Core, and Pol-2 pools eliciting the strongest median responses across all groups (Figure 5B, 5C, S1A). Acute HBV patients had the highest median IFN-γ secretion, while HBV-vaccinated subjects had peak IL-2 responses to the S pool, followed by Core and S-specific responses in acute HBV patients. GrzB secretion was greatest in response to the S pool across all groups. Interestingly, treatment-naïve patients and those who spontaneously resolved CHB exhibited substantially higher GrzB levels than the other groups, suggesting that NUC therapy alters the functional profile of HBV-specific T cells (Figure 5B, 5C S1A, S1B).

The variability in cytokine secretion among CHB patients within the same clinical group precluded the identification of significant differences between groups. In contrast, acute HBV patients had significantly stronger IFN-γ and IL-2 responses to the Core pool, highlighting distinct HBV-specific T cell functions in patients actively clearing the virus (Figure 5D).

### Identification of CHB patients with similar HBV-specific T cell profiles

To identify CHB patients with similar HBV-specific immune responses—based on quantity and combination of secreted cytokines in response to HBV peptide pools— we applied the UMAP algorithm to group samples based on cytokine secretion profiles (IFN-γ, IL-2, GrzB, IL-5; Figure 6A). UMAP showed substantial overlap between IFN-γ and IL-2 secretion but also distinguished samples with high IFN-γ but low IL-2 secretion, or vice versa (Figure 6B). IL-5 secretion was minimal and did not influence clustering, while GrzB secretion formed a distinct cluster. Within this cluster, samples were arranged from low to high GrzB levels, with some overlapping IFN-γ/IL-2 profiles and others showing exclusive GrzB secretion (Figure 6B).

**Fig. 6.**
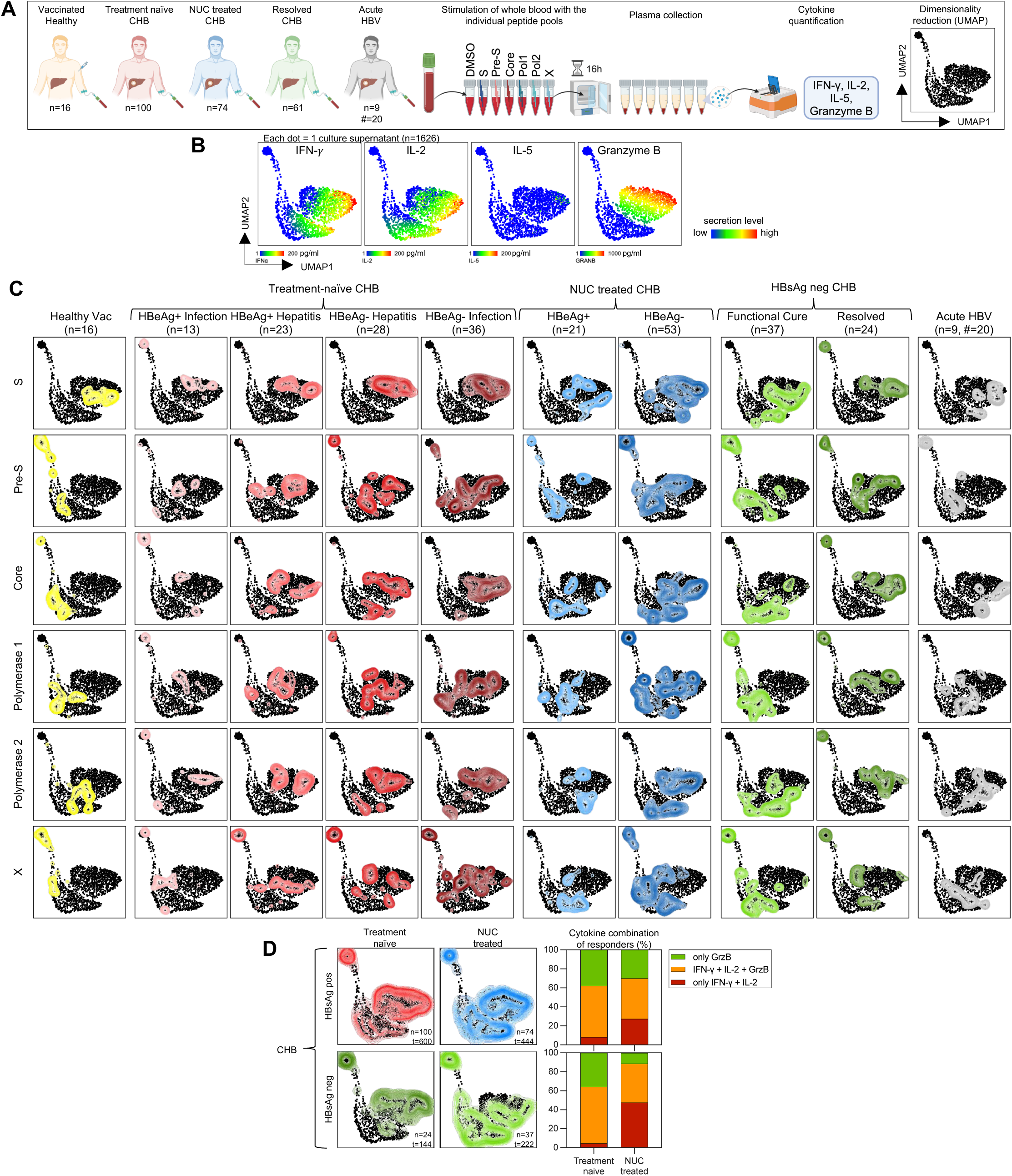
Visualization of HBV-specific T cell secretomes with UMAP. (**A**) Schematic of patients and HBV-CRA. (**B**) UMAP plots generated with all analyzed blood culture supernatants (n=1626) with indicated cytokine secretion heatmaps. (**C**) Concatenated cytokine secretion profiles of peptide-pool stimulated blood from healthy-vaccinated (yellow), different phases of treatment-naïve CHB (shades of red), HBeAg+ and HBeAg- NUC-treated CHB (light and dark blue), HBsAg- CHB (functional cure after NUC, light green; spontaneously resolved, dark green) and patients with acute HBV (HBV-DNA+; grey) overlaid on the global UMAP plot (black dots; each dot corresponds to one culture supernatant). (**D**) Concatenated cytokine secretion profiles of all six peptide-pool stimulated blood samples from HBsAg+ treatment-naïve (n=100; red) and NUC-treated (n=74, blue) CHB patients, and resolved (HBsAg-) treatment-naïve (n=24; dark green) and previously NUC-treated (n=37; light green) CHB patients. Bar graphs (right) show the percentage of blood cultures from treatment-naïve and NUC- treated CHB patients (upper panel: HBsAg+, lower panel: HBsAg-) with cytokine secretion levels >10 pg/mL that contained either only IFN-γ and IL-2 (dark red), IFN-γ, IL-2, and GrzB (orange), or only GrzB (green). n=number of subjects; #=number of samples; t=number of tests.

Figure 6C visualizes the HBV-specific cytokine secretion profiles induced by the six peptide pools in CHB patients, healthy vaccinated controls and acute HBV patients. As expected, in vaccinated controls, all S-peptide pool stimulated samples clustered in regions with medium-to-high IFN-γ, IL-2, and GrzB, while samples stimulated with other pools clustered in the regions with low-to-negative cytokine levels. In contrast, CHB patients of all disease phases exhibited heterogeneous clusters across all peptide pools, with some clustering together (i.e., S in HBeAg-negative hepatitis) and others forming multiple distinct (i.e., S and Core in HBeAg-positive hepatitis, S in HBsAg-negative resolved CHB) or widely dispersed clusters (i.e., responses to all peptide pools in NUC-treated HBeAg-negative CHB), depicting the marked heterogeneity of HBV-specific T cell responses even within the same disease phase. As expected from the analysis in Figure 5, in acute HBV patients, Core peptide pool responses clustered in regions with the highest levels of IFN-γ and IL-2, distributed along a gradient of GrzB secretion ranging from high to none. Moreover, UMAP visualization also revealed differences between treatment-naïve and NUC-treated CHB patients (Figure 6D). In treatment-naïve patients, 92% of responses involved GrzB, either alone (38%) or combined with IFN-γ/IL-2 (54%). In contrast, NUC-suppressed patients had more responses secreting only IFN-γ/IL-2 (27%), a pattern even more pronounced in patients with HBsAg loss. Among spontaneously resolved CHB cases, only 4.6% of responses lacked GrzB, compared to 47.6% in NUC-treated functional cure patients.

Next, to identify and subgroup CHB patients based on their HBV-specific T cell cytokine release profile, we combined the dimensional reduction method UMAP with PhenoGraph, an unsupervised clustering algorithm. With the k-nearest neighbors parameter of 30, PhenoGraph identified 13 distinct subgroups of cytokine release profiles (Figures 7A-C). We analyzed the distribution of samples across the 13 cytokine clusters (Figure 7D). This revealed substantial diversity in T cell responses within the same clinical category. Acute HBV patients predominantly fell into clusters C1–C3, characterized by high IFN-γ and IL-2 secretion, with varying GrzB levels: no (C1), low (C2), or high (C3). In contrast, treatment-naïve CHB patients and those with resolved CHB (spontaneous HBsAg loss) were mostly in C3-C7, marked by high GrzB secretion, either with (C3-C4) or without (C5-C7) IFN-γ and IL-2. Conversely, NUC-suppressed patients and those who achieved functional cure (HBsAg loss after NUCs) showed more responses in C1 and C2, characterized by IFN-γ and IL-2 with no or low GrzB.

**Fig. 7.**
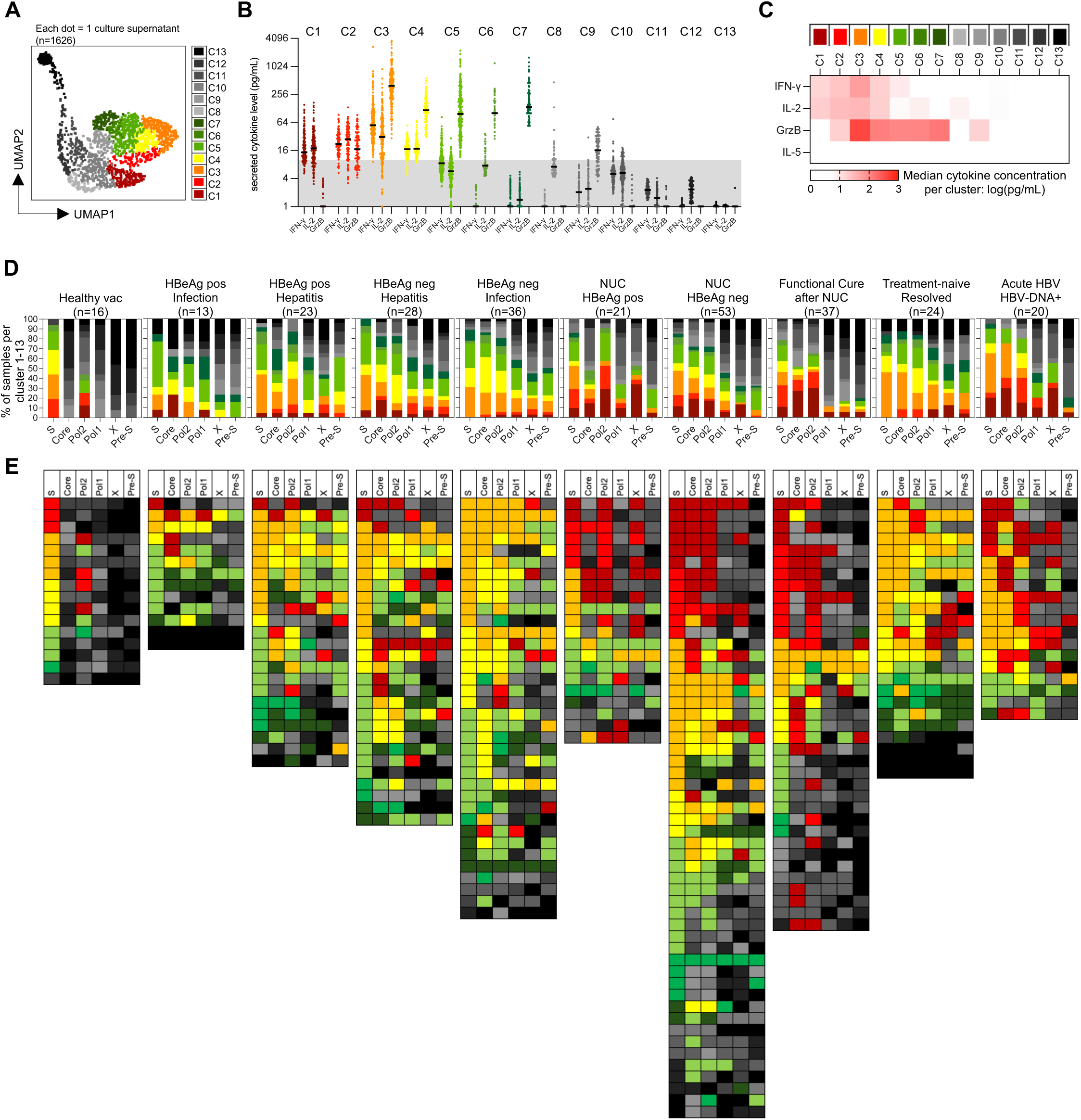
Clustering of samples with similar HBV-specific T cell cytokine secretion profiles with Phenograph. (**A**) UMAP plot with 13 color-coded PhenoGraph clusters. (**B**) Cytokine levels (IFN-γ, IL-2, GrzB) of all samples that fell into the PhenoGraph clusters 1 to 13 (each dot one sample; black line=median). (**C**) Heatmap of median cytokine concentrations of samples in clusters 1 to 13. (**D**) The portions of the 13 clusters identified by PhenoGraph in HBV-CRAs are shown as bar plots for each patient group in response to the six HBV-peptide pools. The colors of the 13 clusters correspond to those of the UMAP in panel (A). (**E**) Each line represents one individual, with the color code indicating the cluster to which their cytokine secretion profile was assigned in response to the HBV peptide pools.

This subgrouping approach allowed us to visualize individual patient response patterns to HBV peptide pools (Figure 7E). While some individuals showed similar cytokine profiles to the six HBV peptide pools, all falling within the same cluster, others exhibited distinct responses to each pool.

### Conclusions

We developed a point-of-care assay, HBV-CRA, to quantitatively and functionally profile T cell immunity against HBV using ∼2.5 mL of whole blood (320 μL per peptide pool for S, Pre-S1/S2, Core, Polymerase, and X). Compatible with high-throughput processing, the assay adapts a method we previously used to measure SARS-CoV-2-specific T cells(23,24), including in low-resource settings(32).

HBV-CRA simplifies testing by using fresh whole blood, eliminating the need for PBMC isolation, which is impractical in most facilities. The two-step process involves 1) overnight blood stimulation with peptide pools and supernatant collection, and 2) cytokine quantification. This modular design enables cytokine analysis requiring specialized equipment to be performed off-site (see ref 32), ensuring standardization and reproducibility. With sensitivity in the 0.01-0.02% HBV-specific T cell range among total T cells, the assay is well-suited to detect the low frequency of HBV-specific T cells typically present in CHB.

Stimulation of CHB patient blood with HBV peptide pools triggered cytokine secretion (primarily IFN-γ, IL-2, and GrzB) in ∼80% of cases, providing a quantitative and qualitative profile of circulating HBV-specific T cells. The HBV-CRA delivered reproducible results in NUC-suppressed patients over three months and exceeded the sensitivity of ex vivo ELISpot assays. In acute HBV, the assay effectively captured the dynamic changes in HBV-specific T cell responses(29,30), albeit arginase modulating T cell function(33). These findings highlight the assay’s utility for monitoring fluctuations of T cell responses that could occur during the initiation or termination of treatment or hepatic flares in CHB patients.

The HBV-CRA offers not only superior sensitivity but also simplifies the assessment of functional HBV-specific T cell heterogeneity. Typically, this analysis would require advanced techniques such as HLA-tetramer staining, intracellular cytokine staining(14), or single-cell transcriptomics. By simultaneously measuring multiple cytokines and combining their concentrations using a multidimensional reduction algorithm, HBV-CRA derives a specific secretome profile for all HBV proteins in each patient. We noted that various secretome profiles could be detected among patients within each clinical phase and that specific secretome profiles did not segregate into any specific clinical category. These findings likely reflect the differential mechanisms of HBV-specific T cell defects observed in CHB(1,17).

Nonetheless, some trends were evident. Patients with acute HBV in the early clearance phase (while still HBV-DNA+) showed high IL-2 and IFN-γ secretion in response to S and Core peptide pools, with relatively low GrzB. Interestingly, this profile was not exclusive to acute HBV patients. We observed similar secretome patterns in some treatment-naïve and NUC-treated CHB patients raising the possibility that patients displaying this “HBV clearance” profile may respond better to specific treatments or maintain HBV control after NUC discontinuation. The protective role of balanced IFN-γ and IL-2 co-production by HBV-specific T cells is supported by a recent study identifying IL-2 as a potential biomarker for viral control and response to checkpoint inhibitors(34).

We also observed that reduced GrzB production by HBV-specific T cells was associated with NUC treatment. A profile characterized by IFN-γ and IL-2 secretion without GrzB was predominant in NUC-treated patients or those who achieved functional cure on NUCs. Conversely, GrzB secretion, alone or in combination with IL-2/IFN-γ, was more frequently observed in treatment-naïve CHB patients, or in those who spontaneously lost HBsAg. This distinction between spontaneous HBsAg loss and NUC-induced HBsAg sero-clearance is intriguing. The reduced GrzB production in NUC-treated patients may reflect the immunomodulatory effects of acyclic nucleoside phosphonates, which can alter monocytes’ cytokine secretion(35). However, the persistence of this profile after treatment cessation warrants further investigation to understand the long-term effects of NUCs on immune responses. These findings underscore the HBV-CRA’s potential to uncover previously unobserved differences in HBV-specific immune responses induced by NUC treatment.

A decade ago, Dammermann et al. described a similar whole-blood CRA for HBV-specific T cell assessment(36). While both assays share a similar approach, our method requires less blood (320 µL vs. 1 mL per stimulation), a shorter time (16 hours vs. 24 hours), and lower peptide concentrations (2 µg/mL vs. 5 µg/mL). Additionally, their study did not compare the HBV-CRA’s sensitivity to classical methods or highlight how it can reveal HBV-specific T cell heterogeneity.

At this juncture, we analyzed IFN-γ, IL-2, GrzB, and IL-5. IFN-γ and IL-2 are critical Th1 cytokines, inhibiting HBV replication(37) and maintaining CD8 T cell fitness(38), respectively. GrzB defines T cells with high lytic capability, and HBV-specific CD8+ T cells with low cytokine production but positive for granzyme and perforin were observed in CHB patients(14). Although IL-5 was undetectable here, we included a Th2 cytokine for its potential relevance in specific patient populations(39). In addition, we already have experimental evidence that therapeutic vaccine-induced HBV-specific T cells can produce Th2 cytokines in some CHB patients(40).

IL-10 and TNF-α were initially included, since IL-10 has been linked to highly functional T cell responses(24) and the promotion of CD8 T cell maturation in HBV transgenic mice(41), while TNF-α has the ability to synergize with IFN-γ to induce noncytolytic HBV-DNA reduction in infected hepatocytes(42), and reports of TNF-α-producing CD4 T cells(43). However, both were excluded from the final analysis due to their fluctuating levels and association with inflammatory states(44) rather than HBV-specific T cell functionality. Future studies should explore additional cytokines like IL-17 and IL-21 to further elucidate HBV-specific T cell responses.

The HBV-CRA has some limitations. It measures only circulating HBV-specific T cells and cannot distinguish whether cytokines originate from CD4+ or CD8+ T cells. Additionally, it cannot detect anergic or functionally exhausted HBV-specific T cells, and using 15-mer peptides may underestimate CD8 T cell frequencies(45). Despite these limitations, the assay offers a minimally invasive and simple approach to obtain both quantitative and qualitative measurements of HBV-specific T cells. It is well-suited for clinical settings, particularly for monitoring CHB patients undergoing novel therapeutic interventions, as we recently demonstrated in our study with siRNA and therapeutic vaccination(40).

In summary, HBV-CRA represents a powerful tool for advancing the understanding and management of chronic HBV. By uncovering functional differences in HBV-specific T cell responses, it provides new insights into the immune dynamics of CHB patients and holds promise for guiding future treatment strategies.

---

**Fig. S1:**
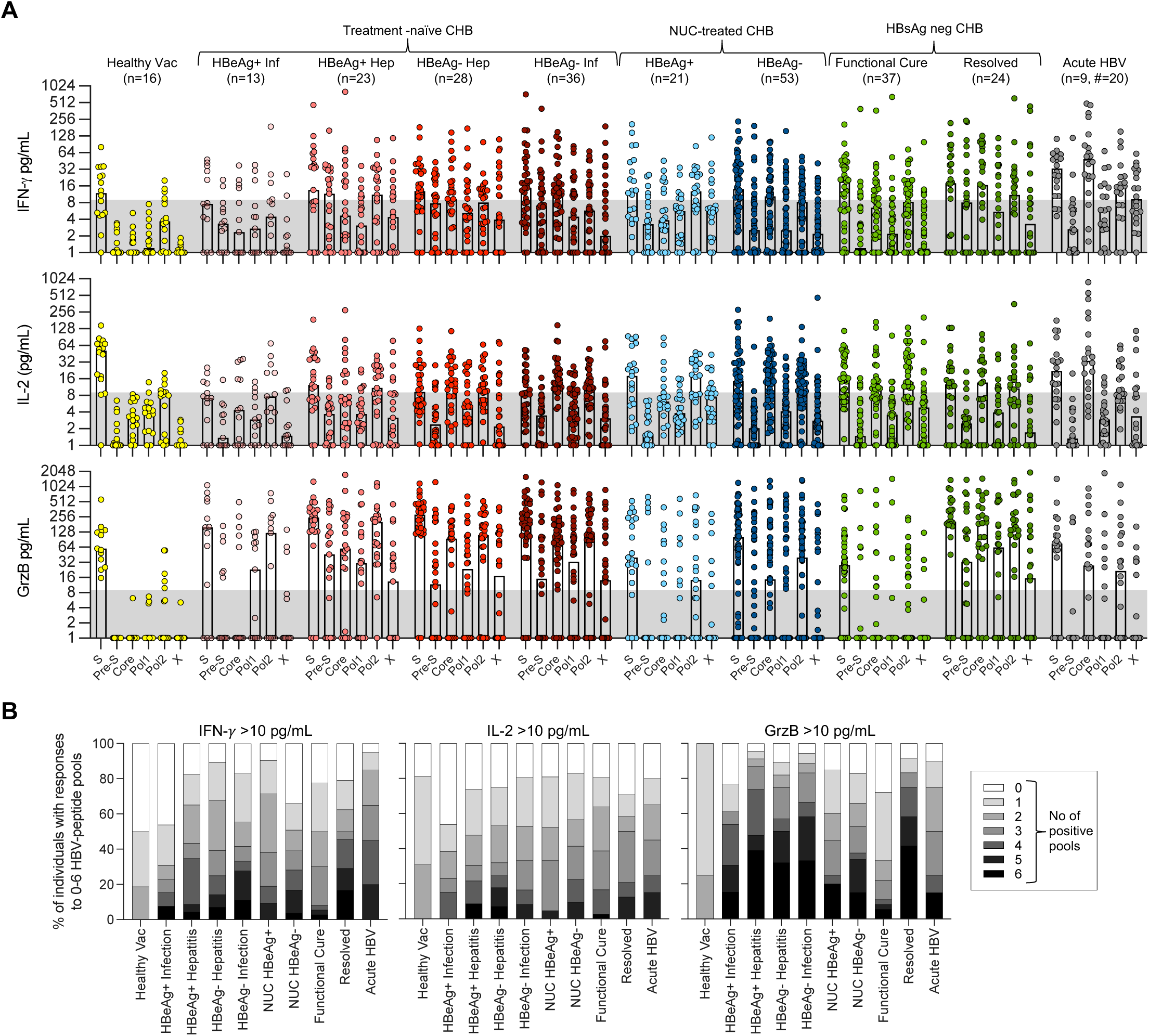
Dominance and multispecificity of HBV-specific T cell responses among patients of different clinical categories. (**A**) Secretion levels (pg/mL) of IFN-γ, IL-2, and GrzB, in response to the indicated HBV-peptide pools are shown for the various patient groups. Each dot represents one patient sample; bar represents median; n=number of subjects; #=number of samples. (**B**) Frequency of samples with cytokine levels >10 pg/mL of IFN-γ, IL-2, and GrzB in response to 0 to 6 different HBV peptide pools.

**Table S1:**
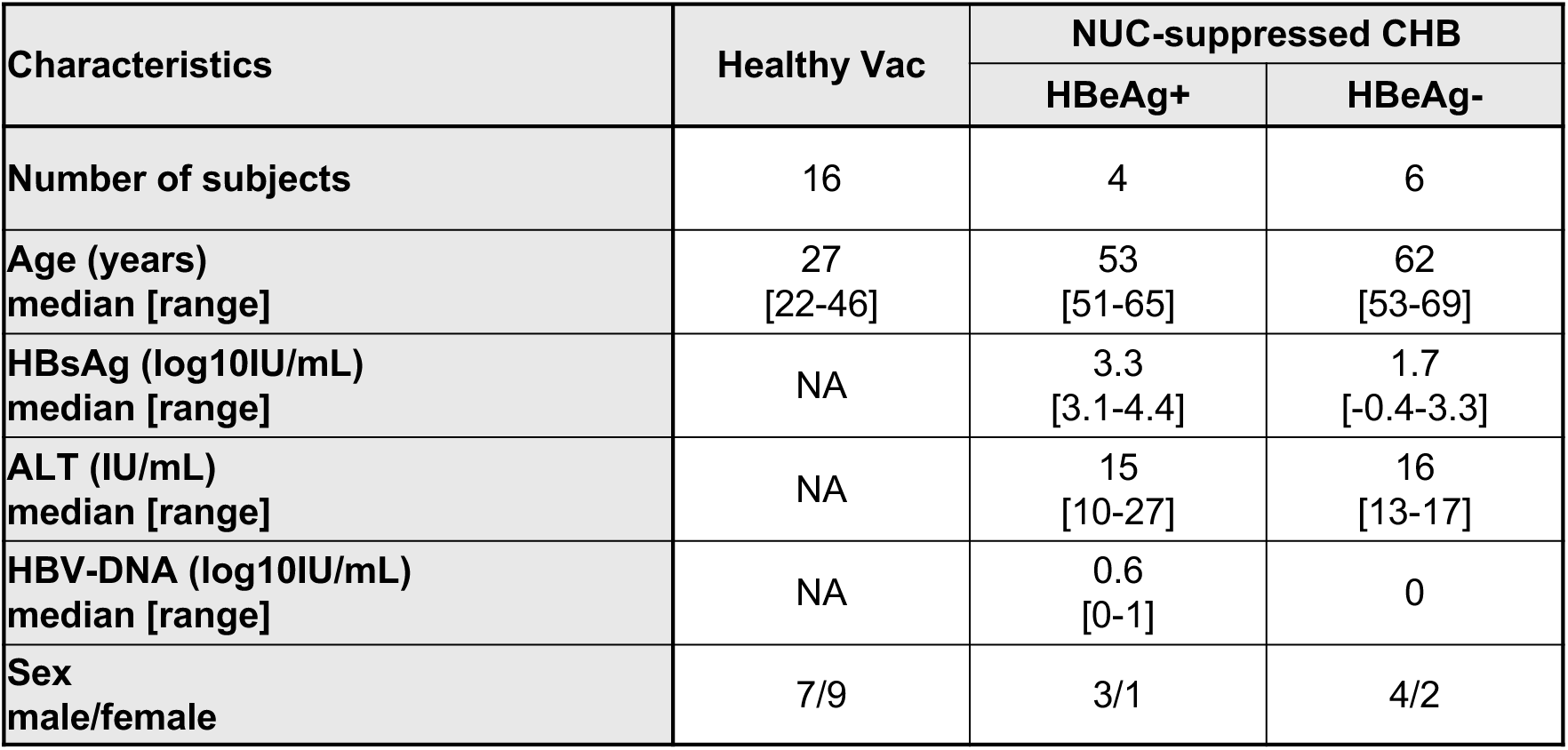
Clinical and virological characteristics of healthy subjects and NUC-suppressed CHB patients followed monthly.

**Table S2:**
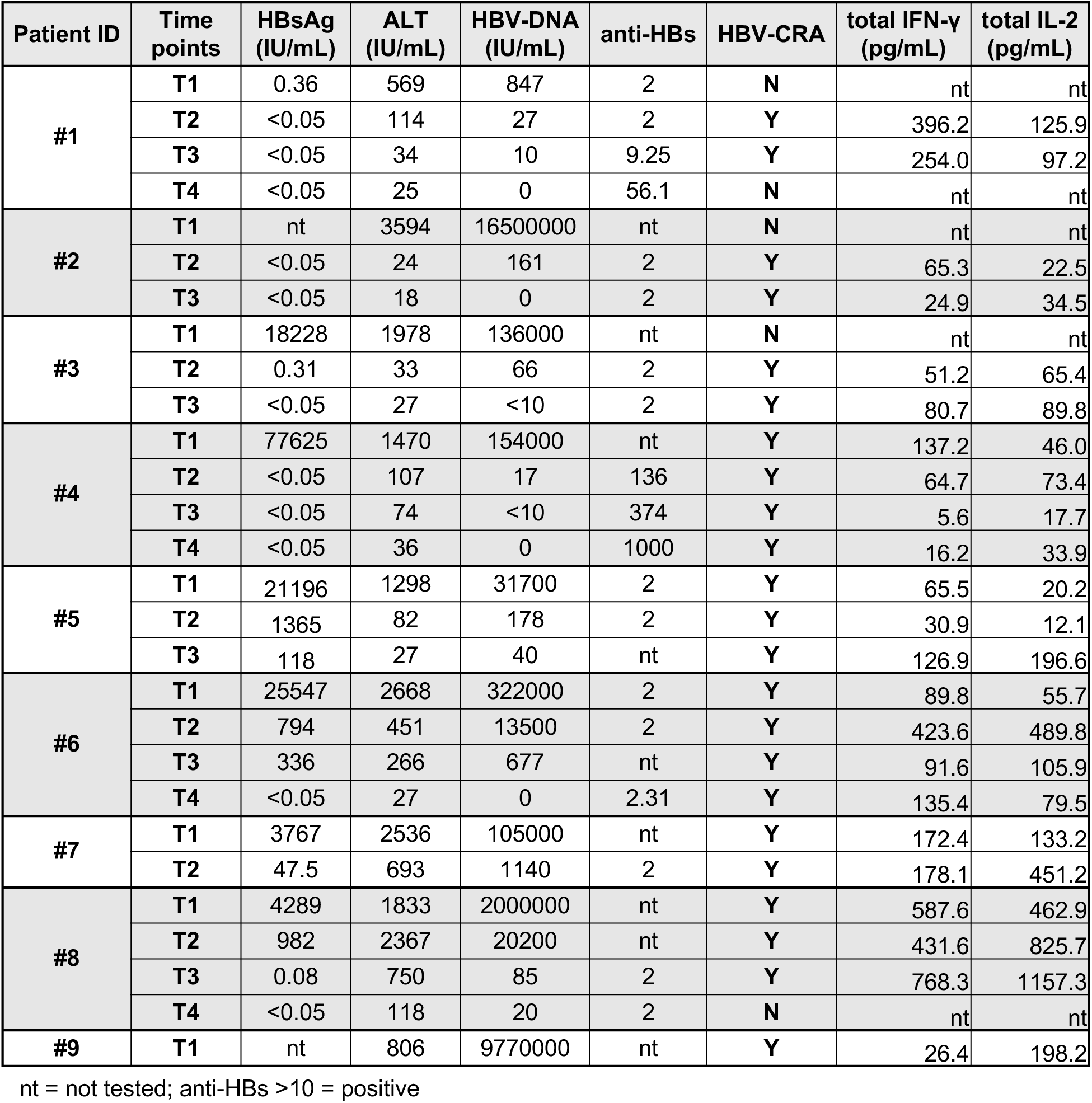
Clinical and virological characteristics of acute HBV patients in this study. HBV-CRA was performed on indicated samples. The sum of IFN-γ and IL-2 secreted in response to the six HBV-peptide pools (Core, S, PreS, Pol-1, Pol-2, X) is shown.

**Table S3:**
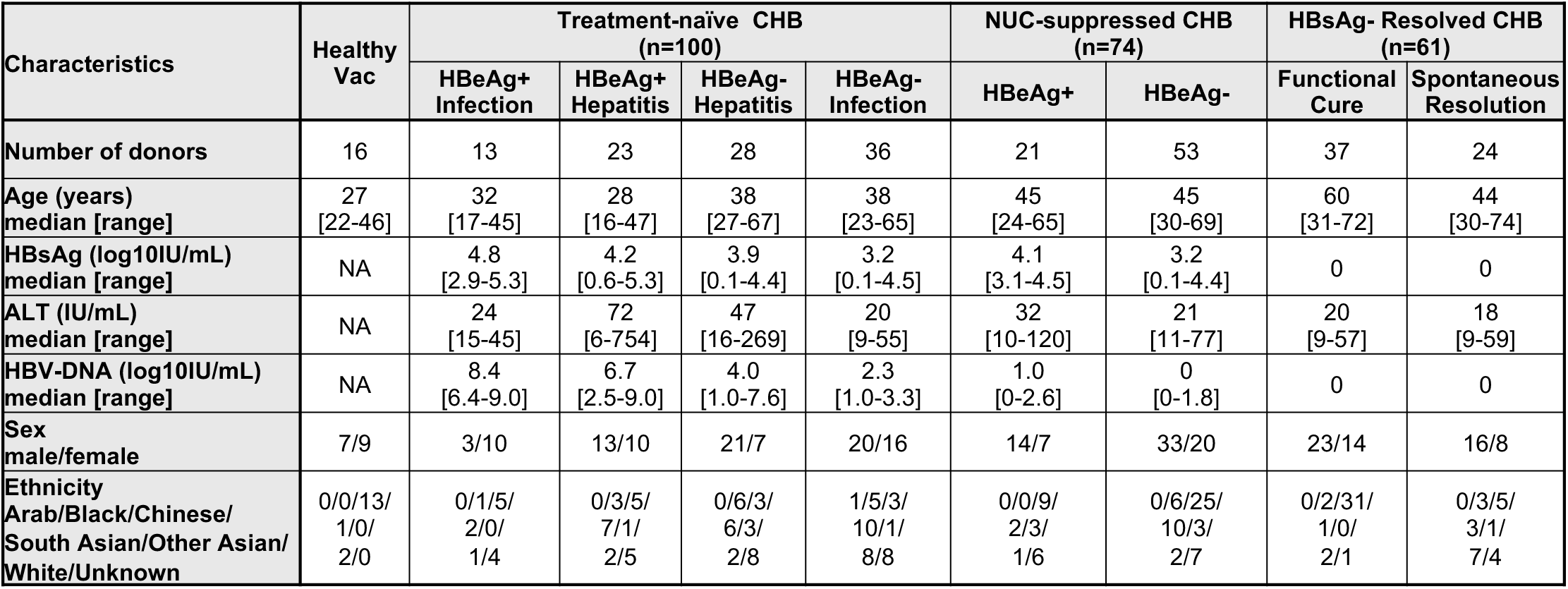
Clinical and virological characteristics of all CHB patients in this study.

