## Supplementary material for "Heterogeneous T cell responses across Hepatitis B virus clinical phases revealed by rapid whole-blood HBV T cell analysis": Supplemantal Tables S1-3, and Figure S1

**Table S1: Clinical and virological characteristics of healthy subjects and NUC-suppressed CHB patients followed monthly**

| Characteristics | Healthy Vac | NUC-suppressed CHB | |
|---|---|---|---|
| | | HBeAg+ | HBeAg- |
| Number of subjects | 16 | 4 | 6 |
| Age (years)<br>median [range] | 27<br>[22-46] | 53<br>[51-65] | 62<br>[53-69] |
| HBsAg (log10IU/mL)<br>median [range] | NA | 3.3<br>[3.1-4.4] | 1.7<br>[-0.4-3.3] |
| ALT (IU/mL)<br>median [range] | NA | 15<br>[10-27] | 16<br>[13-17] |
| HBV-DNA (log10IU/mL)<br>median [range] | NA | 0.6<br>[0-1] | 0 |
| Sex<br>male/female | 7/9 | 3/1 | 4/2 |

**Table S2: Clinical and virological characteristics of acute HBV patients in this study**

HBV-CRA was performed on indicated samples. The sum of IFN-γ and IL-2 secreted in response to the six HBV-peptide pools (Core, S, PreS, Pol-1, Pol-2, X) is shown.

| Patient ID | Time points | HBsAg (IU/mL) | ALT (IU/mL) | HBV-DNA (IU/mL) | anti-HBs | HBV-CRA | total IFN-γ (pg/mL) | total IL-2 (pg/mL) |
|---|---|---|---|---|---|---|---|---|
| #1 | T1 | 0.36 | 569 | 847 | 2 | N | nt | nt |
| | T2 | <0.05 | 114 | 27 | 2 | Y | 396.2 | 125.9 |
| | T3 | <0.05 | 34 | 10 | 9.25 | Y | 254.0 | 97.2 |
| | T4 | <0.05 | 25 | 0 | 56.1 | N | nt | nt |
| #2 | T1 | nt | 3594 | 16500000 | nt | N | nt | nt |
| | T2 | <0.05 | 24 | 161 | 2 | Y | 65.3 | 22.5 |
| | T3 | <0.05 | 18 | 0 | 2 | Y | 24.9 | 34.5 |
| #3 | T1 | 18228 | 1978 | 136000 | nt | N | nt | nt |
| | T2 | 0.31 | 33 | 66 | 2 | Y | 51.2 | 65.4 |
| | T3 | <0.05 | 27 | <10 | 2 | Y | 80.7 | 89.8 |
| #4 | T1 | 77625 | 1470 | 154000 | nt | Y | 137.2 | 46.0 |
| | T2 | <0.05 | 107 | 17 | 136 | Y | 64.7 | 73.4 |
| | T3 | <0.05 | 74 | <10 | 374 | Y | 5.6 | 17.7 |
| | T4 | <0.05 | 36 | 0 | 1000 | Y | 16.2 | 33.9 |
| #5 | T1 | 21196 | 1298 | 31700 | 2 | Y | 65.5 | 20.2 |
| | T2 | 1365 | 82 | 178 | 2 | Y | 30.9 | 12.1 |
| | T3 | 118 | 27 | 40 | nt | Y | 126.9 | 196.6 |
| #6 | T1 | 25547 | 2668 | 322000 | 2 | Y | 89.8 | 55.7 |
| | T2 | 794 | 451 | 13500 | 2 | Y | 423.6 | 489.8 |
| | T3 | 336 | 266 | 677 | nt | Y | 91.6 | 105.9 |
| | T4 | <0.05 | 27 | 0 | 2.31 | Y | 135.4 | 79.5 |
| #7 | T1 | 3767 | 2536 | 105000 | nt | Y | 172.4 | 133.2 |
| | T2 | 47.5 | 693 | 1140 | 2 | Y | 178.1 | 451.2 |
| #8 | T1 | 4289 | 1833 | 2000000 | nt | Y | 587.6 | 462.9 |
| | T2 | 982 | 2367 | 20200 | nt | Y | 431.6 | 825.7 |
| | T3 | 0.08 | 750 | 85 | 2 | Y | 768.3 | 1157.3 |
| | T4 | <0.05 | 118 | 20 | 2 | N | nt | nt |
| #9 | T1 | nt | 806 | 9770000 | nt | Y | 26.4 | 198.2 |

nt = not tested; anti-HBs >10 = positive

### Table S3: Clinical and virological characteristics of all CHB patients in this study

| Characteristics | Healthy Vac | Treatment-naïve CHB (n=100) | | | | NUC-suppressed CHB (n=74) | | HBsAg- Resolved CHB (n=61) | |
|---|---|---|---|---|---|---|---|---|---|
| | | HBeAg+ Infection | HBeAg+ Hepatitis | HBeAg- Hepatitis | HBeAg- Infection | HBeAg+ | HBeAg- | Functional Cure | Spontaneous Resolution |
| **Number of donors** | 16 | 13 | 23 | 28 | 36 | 21 | 53 | 37 | 24 |
| **Age (years) median [range]** | 27 [22-46] | 32 [17-45] | 28 [16-47] | 38 [27-67] | 38 [23-65] | 45 [24-65] | 45 [30-69] | 60 [31-72] | 44 [30-74] |
| **HBsAg (log10IU/mL) median [range]** | NA | 4.8 [2.9-5.3] | 4.2 [0.6-5.3] | 3.9 [0.1-4.4] | 3.2 [0.1-4.5] | 4.1 [3.1-4.5] | 3.2 [0.1-4.4] | 0 | 0 |
| **ALT (IU/mL) median [range]** | NA | 24 [15-45] | 72 [6-754] | 47 [16-269] | 20 [9-55] | 32 [10-120] | 21 [11-77] | 20 [9-57] | 18 [9-59] |
| **HBV-DNA (log10IU/mL) median [range]** | NA | 8.4 [6.4-9.0] | 6.7 [2.5-9.0] | 4.0 [1.0-7.6] | 2.3 [1.0-3.3] | 1.0 [0-2.6] | 0 [0-1.8] | 0 | 0 |
| **Sex male/female** | 7/9 | 3/10 | 13/10 | 21/7 | 20/16 | 14/7 | 33/20 | 23/14 | 16/8 |
| **Ethnicity Arab/Black/Chinese/ South Asian/Other Asian/ White/Unknown** | 0/0/13/ 1/0/ 2/0 | 0/1/5/ 2/0/ 1/4 | 0/3/5/ 7/1/ 2/5 | 0/6/3/ 6/3/ 2/8 | 1/5/3/ 10/1/ 8/8 | 0/0/9/ 2/3/ 1/6 | 0/6/25/ 10/3/ 2/7 | 0/2/31/ 1/0/ 2/1 | 0/3/5/ 3/1/ 7/4 |

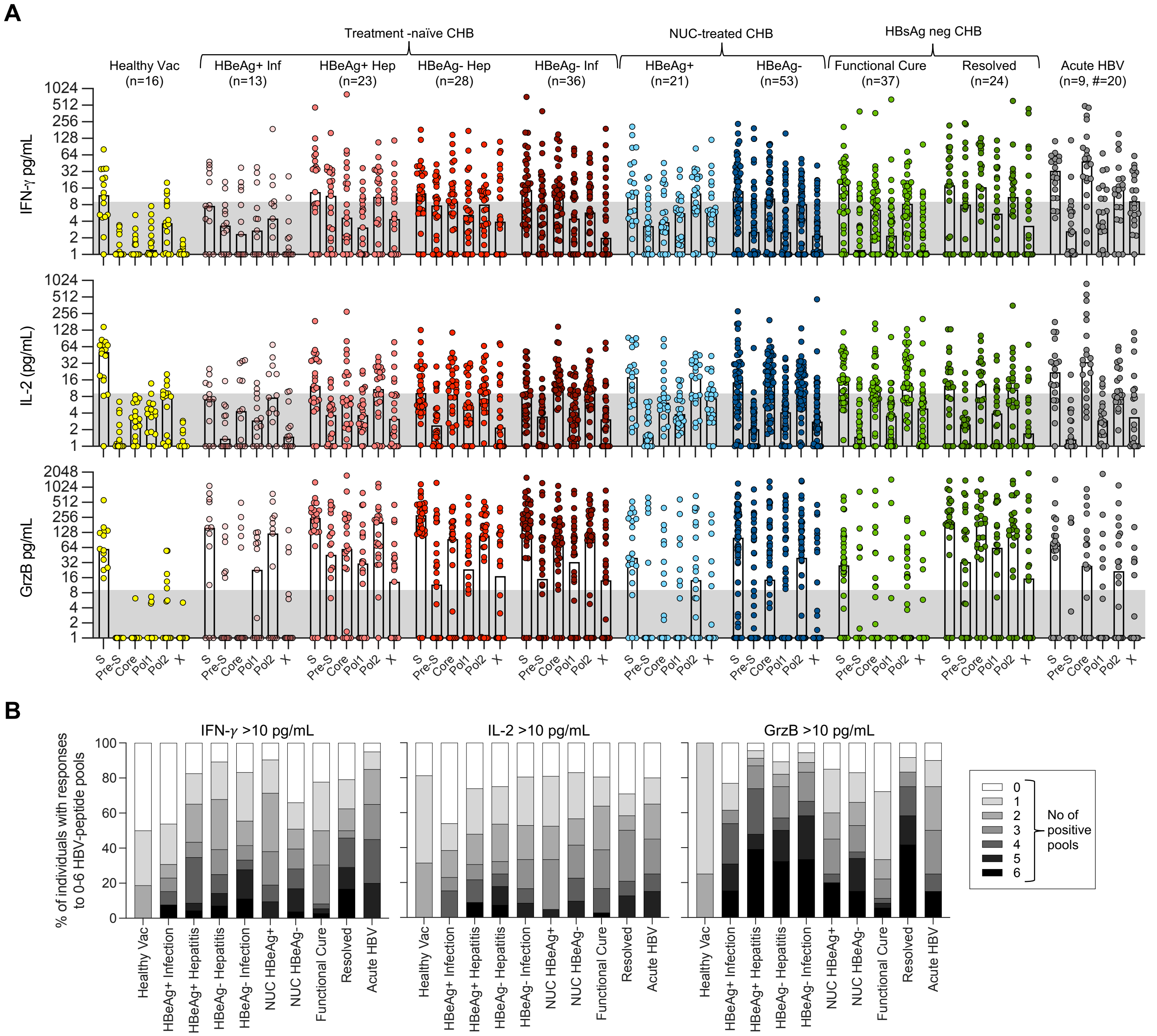


**Fig. S1: Dominance and multispecificity of HBV-specific T cell responses among patients of different clinical categories**

(**A**) Secretion levels (pg/mL) of IFN-γ, IL-2, and GrzB, in response to the indicated HBV-peptide pools are shown for the various patient groups. Each dot represents one patient sample; bar represents median; n=number of subjects; #=number of samples. (**B**) Frequency of samples with cytokine levels >10 pg/mL of IFN-γ, IL-2, and GrzB in response to 0 to 6 different HBV peptide pools.
